# Distribution-Constrained Optimization for Reliable ML-Guided 5′UTR Sequence Design

**DOI:** 10.64898/2026.08.03.742388

**Authors:** Ryohei Yamaguchi, Chinatsu Mori, Sota Inoue

## Abstract

The 5′ untranslated region (5′ UTR) shapes translation initiation, so its design is central to mRNA therapeutics and to improving protein-production cell lines. Deep-learning models that predict translation efficiency, measured as mean ribosome load (MRL), from the 5′ UTR sequence have been combined with genetic algorithms (GAs) for sequence optimization. However, optimizing against a model trained on offline data risks reward hacking that exploits the model’s estimation error outside the training distribution, yielding sequences that score highly in prediction yet fail to perform in the wet lab. Yet for 5′ UTR design, few studies have systematically examined which region should be treated as untrustworthy (the definition of out-of-distribution, OOD) or which constraints keep the search away from it.

We present a constrained optimization that keeps candidates within a trust region where the predictor’s validated accuracy holds; here “reliable” denotes keeping candidates within the training distribution over which prediction has been validated, not a guarantee of measured performance. As the OOD score, we compare the k-nearest-neighbor (KNN) distance in the predictor’s embedding space against a pseudo-perplexity (PPPL) from the encoder and LM head, and show that for nucleotide sequences—whose vocabulary is small—PPPL fails to separate in- vs out-of-distribution, whereas the KNN distance is an effective OOD score that can define a trust region even from unlabeled native UTR sequences. Using the KNN distance as a hard GA constraint keeps all candidates inside the trust region while maintaining predicted MRL: under unconstrained optimization most final-generation candidates (72–96% across seeds) left the trust region (self-KNN p95), whereas the hard constraint holds predicted MRL at the unconstrained level and yields about 4.3*×* more selectable low-risk candidates than post-hoc filtering of the unconstrained output. Comparing an output extrapolation guard, reference-sequence similarity and structural accessibility (RNAplfold), we find that the guard and the similarity constraint also suppress OOD as a side effect, whereas making accessibility a secondary objective broadens the search without suppressing OOD.

## Introduction

The 5′ UTR is a key determinant of mRNA translation efficiency and contributes substantially to protein productivity. Methods for predicting translation efficiency from sequence and for optimizing the sequence have therefore been developed.

Bhandari et al. and Terai et al. estimated accessibility from the base-pairing probabilities around the translation start site of the mRNA and showed that it is useful for estimating translation efficiency (Terai and Asai, 2020; Bhandari et al., 2019, 2021). Sample et al. (2019) proposed Optimus 5-Prime, a deep-learning model that predicts MRL, the average number of ribosomes loaded on an mRNA. Subsequent MRL predictors advanced to UTR-Insight (Pan et al., 2025), which combines a Frame-pool block that handles arbitrary lengths with a ConvTransformer that captures both short- and long-range information. Genetic algorithms are widely used for sequence design that uses these models as evaluators (Sample et al., 2019; Pan et al., 2025).

However, optimization that uses a predictor trained on offline data as the objective risks reward hacking that exploits the estimation error outside the training distribution, producing sequences that score highly in prediction but fail to perform experimentally (Kim et al., 2025). This failure of the predictor in its extrapolation regime has also been reported in other ML-guided sequence design tasks, such as protein design (Brookes et al., 2019). Yet few studies have systematically evaluated, for 5′ UTR design, which region to regard as untrustworthy (the definition of OOD) or which constraints keep the search away from it.

We therefore present a constrained optimization toward a trust region (Fig. 1). Specifically, we show that (i) the KNN distance in the predictor’s embedding space serves as an OOD score (established by comparing its discriminative behavior with that of the pseudo-perplexity PPPL), so that a trust region can be defined even from unlabeled native UTR sequences alone; (ii) using it as a hard constraint keeps low-risk candidates inside the trust region without sacrificing predicted MRL; and (iii) several search-space controls, imposed as constraints or objectives, affect the reliability and diversity of the candidate distribution.

**Figure 1.**
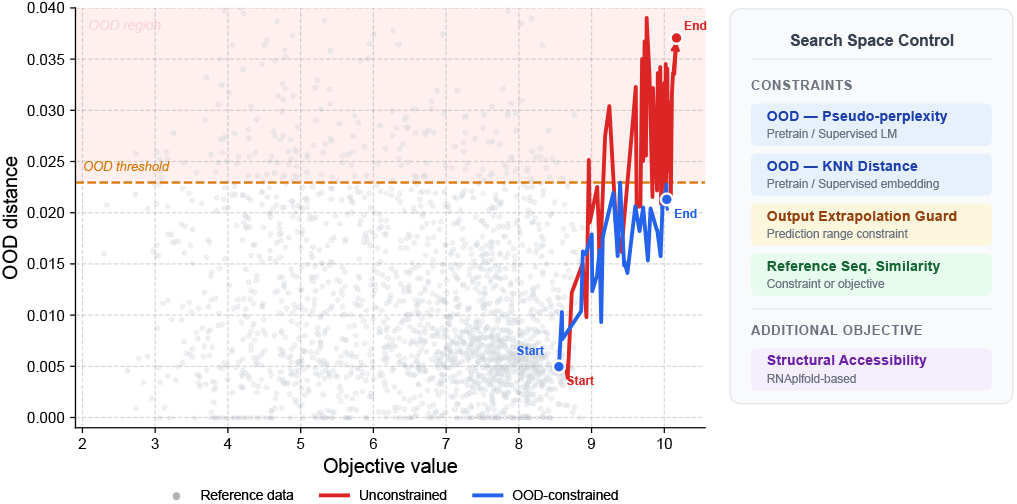
Conceptual overview of OOD departure in ML-guided optimization and of search-space control by constraints. Optimization trajectories in the space of predicted MRL and KNN distance. Grey points show the self-KNN distance distribution of the pretraining data, and the dashed line marks the p90 threshold. Whereas the unconstrained run departs into the OOD region, the OOD-constrained run stays within the threshold. The search-space controls introduced in this study are also listed.

## Methods

### Sequence optimization

For in silico design of 5′ UTR sequences, we used the deep-learning model UTR-Insight (Pan et al., 2025), which predicts translation efficiency from sequence, as the evaluator, and performed sequence optimization with a genetic algorithm. We used NSGA-II (pymoo 0.6.1.5) with a population size of 256 for 1,000 generations at seed=42. The three main conditions (Base, KNN-Pr, KNN-Sv) were checked for reproducibility across three seeds (seed 42–44); unless otherwise noted, quantitative values are based on the final generation (generation 1000) of seed 42. Sequences were represented by integer encoding ({0,1,2,3}={A,C,G,U}, 60 variable positions), and the initial population was sampled uniformly at random at each position. Crossover used an adaptive scheme (two-point crossover in even generations, uniform crossover in odd generations, probability 0.9). Mutation was applied independently to each base, with its probability decayed linearly from 0.05 to 0.025 over generations. Selection used a binary tournament (non-dominated rank plus crowding distance), and duplicate removal was enabled. After crossover and mutation in each generation, a repair operator removed internal AUGs, broke homopolymer runs longer than five bases, and restored the fixed motifs (the 3′-end Kozak sequence and a splicing motif). UTR-Insight was implemented in-house in code based on ESM2 (Lin et al., 2023) using PyTorch 2.0.1. Its high agreement with the test-set predictions reported in the original paper (Pan et al., 2025) (R*≈*1.00) confirms the validity of the reimplementation.

### Search space control

In addition to the GA optimization that uses the predicted translation efficiency as the objective, we designed four indicators to control the search space.

### OOD constraints

Perplexity is an evaluation metric that expresses how well a language model can predict a sequence (Jurafsky and Martin”, 2026); here we used a pseudo-perplexity, PPPL, computed from the encoder and LM head of UTR-Insight as an OOD score, defined as

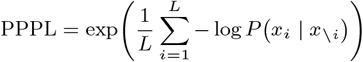

That is, by masking each base one at a time we compute the conditional probability; the higher the PPPL, the more unnatural the sequence is to the model and the less its prediction can be trusted. We also used the KNN distance to the training data as an OOD score. The embeddings were obtained by mean-pooling the last-layer hidden representations of the UTR-Insight encoder over non-padding positions and applying L2 normalization. We used a neighbor count of k=5 and the cosine distance. This follows the standard practice of KNN-based OOD detection (Sun et al., 2022); the setting of that method, which applies the Euclidean distance after L2 normalization, is monotonically equivalent to our cosine distance (identical neighbor ranking) between normalized vectors through the relation 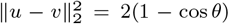. The operational OOD threshold was the percentile computed from the self-KNN distance distribution of the training data itself. As the target training data, we used the corpus used for pretraining (Chu et al., 2024; Cao et al., 2021; Cunningham et al., 2021) and the corpus used for supervised training (Sample et al., 2019), and compared the difference in space control between the two distinct references. The pretraining data consist of five Ensembl species + Cao et al. (255,795 sequences) and do not include the dataset of Sample et al. used for supervised training.

### Output extrapolation guard

A genetic algorithm risks fitting outside the range of the objective’s training-data distribution and selecting sequences whose prediction error is large and whose translation efficiency is not in fact high. To address this, we computed percentiles from the MRL distribution of the GSE114002 dataset (432,563 sequences) (Sample et al., 2019) used for supervised training, and treated a candidate as violating the constraint when its predicted MRL exceeded this threshold (hereafter, the OutEx constraint). Specifically, for the predicted MRL *ŷ* of a candidate we defined the constraint-violation amount as

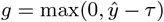

where *τ* is a percentile of the training-data MRL distribution, and *g >* 0 marks a constraint violation (an infeasible solution). Here we used p95 (*τ* = 8.28) as the main condition.

### Structural accessibility

Because secondary-structure formation around the translation start site hinders the binding and scanning of the 40S ribosomal subunit, the accessibility of this region is known to govern translation efficiency (Kozak, 1989). We therefore used the RNAplfold algorithm of ViennaRNA 2.7.2 to compute an accessibility based on base-pairing probabilities, introduced it as a secondary objective of the optimization, and explored its trade-off with predicted MRL.

We concatenated the CDS 5′ start sequence to the 5′ UTR sequence and evaluated the structural openness near the start codon. The RNAplfold parameters were a folding window size *W* = 120, a maximum base-pair span *L* = 120, and a maximum unpaired region length *u* = 30. For the 30 positions comprising 15 nt upstream and 15 nt downstream of the start codon, we computed the probability *p*_unpaired_(*i*) that a single base is unpaired at each position, and took its mean *S*_acc_ as the accessibility score.

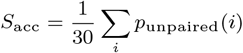

### Reference sequence similarity

Liu et al. (2025) *point out that conventional 5*′ UTR design methods aim to search for broadly applicable sequences and do not consider the sequence of a specific UTR. Other studies also report that the appropriate 5′ UTR differs by CDS gene (Chen et al., 2022). To address this, we introduced the similarity to a reference sequence as an indicator and observed how the search space changed when it was added as a secondary objective and when it was placed in the constraints. The similarity was computed as the position-aligned identity between the candidate and reference sequences,

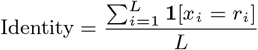

where *x*_*i*_ is the candidate sequence, *r*_*i*_ is the *i*-th base of the reference sequence, and *L* is the length of the shorter of the two sequences. For this identity we set a minimum threshold for a candidate to be feasible,

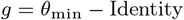

and took *g ≤* 0 (i.e. Identity *≥ θ*_min_) as the constraint. Here we used *θ*_min_ = 0.80.

### Evaluation of candidate space

To characterize the candidate sequences obtained by optimization, we used the following visualizations.

Distribution comparison (violin plots): for the candidate sequences of each optimization condition at the final generation (generation 1000), we drew violin plots of the distributions of three indicators— predicted MRL, the KNN distance to the pretraining data, and the KNN distance to the supervised-training data. Kernel density estimation used a Gaussian kernel with Scott’s rule for bandwidth selection, and the interquartile range and median were overlaid as a box.

KNN-distance scatter plot: we plotted the KNN distance to the pretraining data (horizontal axis) against the KNN distance to the supervised-training data (vertical axis) to visualize the correlation of the OOD characteristics with respect to the two references. The p95 threshold of each reference was shown as a reference line.

UMAP embedding: to visualize the candidate distribution in a low-dimensional space, we performed dimensionality reduction with UMAP (McInnes et al., 2018). We used, as features, 128-dimensional vectors obtained by mean-pooling the last-layer hidden representations of the UTR-Insight encoder (ESM2-based, 6 layers, 128 dimensions) over non-padding positions and applying L2 normalization. After first compressing to 50 dimensions with PCA, we projected to two dimensions with UMAP (neighbor count *k* = 15, minimum distance = 0.1, cosine distance). The PCA and UMAP models were fitted on the combined set of the pretraining data (255,795 sequences) and the supervised-training data (432,563 sequences), and the candidate sequences of each optimization condition were projected (transformed) onto this fixed coordinate system, enabling comparison in a coordinate system consistent across all conditions.

## Results

### OOD Score Selection

We examined which of two scores has the higher ability to capture out-of-distribution (OOD) data: PPPL, which expresses the naturalness to the model using the encoder and LM head of the deep-learning model UTR-Insight, and the KNN distance to the training data in the model’s embedding space. For OOD detection, we assumed setting the 90/95/99 percentiles of each score on the training data as thresholds. We performed GA sequence optimization targeting the predicted value of MRL (Mean Ribosome Load), a measured index of translation efficiency. For the KNN distance, the fraction of final-generation individuals exceeding the p95 threshold reached 72–96% (median 76.6%) across all three seeds (seed 42–44), showing that risky sequences can be quantified (p90 exceedance 97.7% and p99 exceedance 1.6% are seed-42 reference values). The KNN distance also showed an increasing trend, rising 4.7-fold in the median from the first generation, supporting the hypothesis that ML-guided GA optimization carries a latent risk. For PPPL, in contrast, 100% of solutions in all generations were within p90, revealing that PPPL is uninformative for this problem. This suggests that because the nucleotide alphabet has only four choices (A, C, G, U)—extremely small compared with natural language—the training-data perplexity distribution covers almost any input. On the basis of these results, we adopted the KNN distance in the embedding space as the OOD score. Note that this comparison is not an evaluation of discriminative performance against a labeled OOD set, but a contrast of the two scores in their suitability as indicators for capturing the distributional departure that arises during the optimization loop.

### Search Space Control

#### OOD constraints

Considering the model-hacking risk of offline GA optimization, we introduced the OOD score as a hard constraint. Violin plots of the predicted MRL and OOD score with and without the constraint (Baseline) are shown in Figure 2. KNN-Pr denotes the KNN distance to the pretraining data and KNN-Sv the KNN distance to the data used for supervised training. KNN-Pr references the pretraining corpus of native UTR sequences, whereas KNN-Sv references the synthetic MPRA library used for supervised fine-tuning; the two references thus correspond to qualitatively different sequence distributions (natural vs. designed).

**Figure 2.**
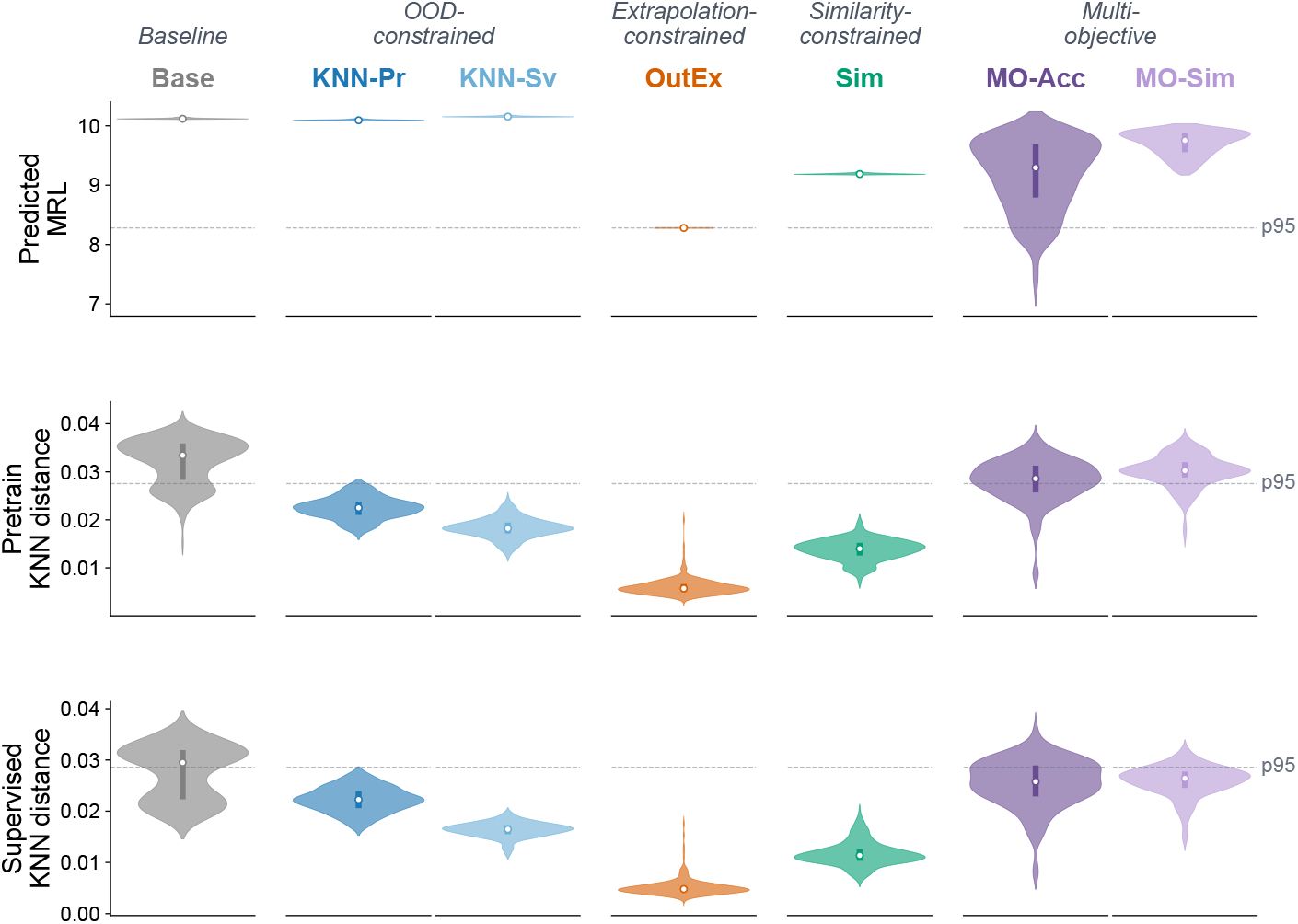
Distribution comparison of the candidate sequences at the final generation (generation 1000) for each optimization condition. Top: predicted MRL; middle: KNN distance to the pretraining data (KNN-Pr); bottom: KNN distance to the supervised-training data (KNN-Sv). Dashed lines mark the p95 threshold of each indicator. Conditions, from left: Base (unconstrained), KNN-Pr/KNN-Sv (OOD constraints), OutEx (output extrapolation guard), Sim (similarity constraint), MO-Acc (accessibility secondary objective), and MO-Sim (similarity secondary objective).

For the Baseline, most predicted MRLs are high, above 10, but the OOD score has a bimodal distribution, one mode of which exceeds the p95 threshold for both the pretraining and the supervised-training data. When the OOD constraint was imposed, the predicted-MRL distribution was kept at the same level as the Baseline while all individuals fell within the OOD-score threshold. This behavior was reproduced across three seeds (seed 42–44): under the KNN-Pr constraint, the median predicted MRL was maintained at the Baseline level (9.8–10.1, variation *±*1.5%) for all seeds. Even a constraint that references only the unlabeled native sequences (Pr) robustly keeps candidates within the trust region while preserving predicted MRL.

The UMAP (Fig. 3) shows that the OOD-constrained optimization did not explore part of the space explored by the Baseline, and that its search extended into a different space; ultimately it produced solutions from a low-risk, high-MRL space.

**Figure 3.**
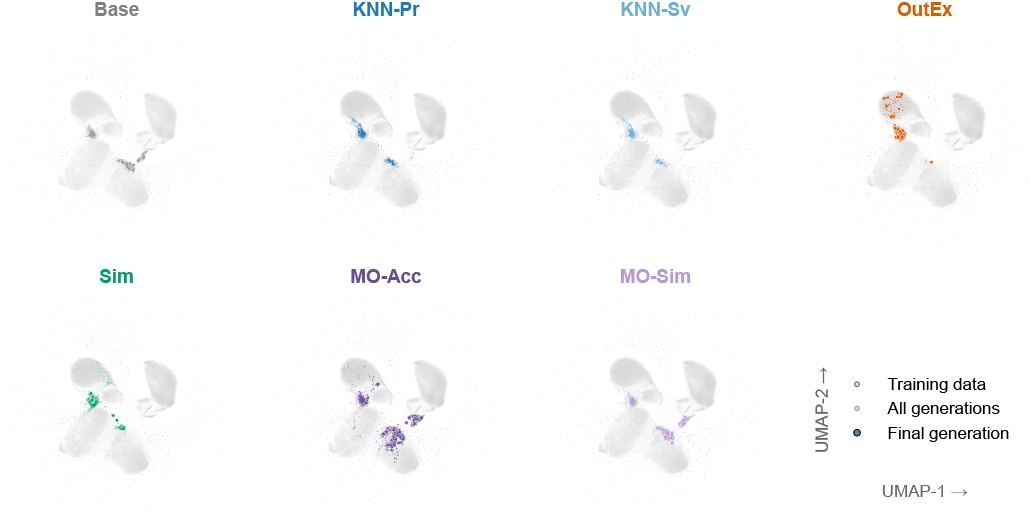
Visualization of the candidate distribution by UMAP embedding. Each panel shows a different optimization condition. Grey points are the training data (pretraining + supervised training) and colored points are the candidate sequences of each condition. Darker points are the final generation (generation 1000).

Quantitatively, the fraction of Baseline individuals falling within the trust region of both references is only 23.4% in the median (3.9–27.7% across seeds). That all individuals fall within the region under the constraint is a consequence of the constraint’s definition; the significance of our method lies in (i) the baseline risk rate— that most candidates depart under no constraint—and (ii) that this departure can be resolved without sacrificing predicted MRL. Relative to a strategy that post-hoc filters the unconstrained output under the same computational budget, the number of selectable low-risk candidates is about 4.3*×* in the median.

Figure 4 shows the difference in scores due to the difference in training data used for the OOD computation. Pr is a native sequence dataset and Sv is the synthetic-sequence dataset of Sample et al.; although computed from qualitatively different datasets, they had a positive Pearson correlation of 0.94 (Base 0.96 / KNN-Pr 0.80 / KNN-Sv 0.83). Individuals obtained by optimizing under the synthetic-sequence set Sv as the constraint tended to show smaller scores than those obtained under Pr. Meanwhile, the point cloud was systematically shifted below the y=x diagonal, so that for the same candidate the KNN-Pr distance tended to exceed the KNN-Sv distance. Dividing into four quadrants at the p95 threshold of each reference, the (inside Pr, outside Sv) quadrant is empty for all conditions and all seeds, whereas the (outside Pr, inside Sv) quadrant is occupied (22–55% for Base). That is, the trust region of Pr is operationally contained in that of Sv. Consequently, the KNN-Pr constraint simultaneously satisfies the trust regions of both references (100% reproduced across three seeds), whereas the KNN-Sv constraint guarantees only the Sv side and the Pr side was seed-dependent (at seed 44, 75.4% of candidates exceeded the Pr-side threshold). This asymmetry reinforces the advantage of using the label-free native-sequence reference (KNN-Pr). We also checked the arrangement of the two references in low-dimensional projections: in the two-dimensional PCA (variance explained by PC1 and PC2 of 53% and 16%) the two distributions overlapped greatly, whereas in UMAP they were largely separated, so the appearance differed between the two 2D projections.

**Figure 4.**
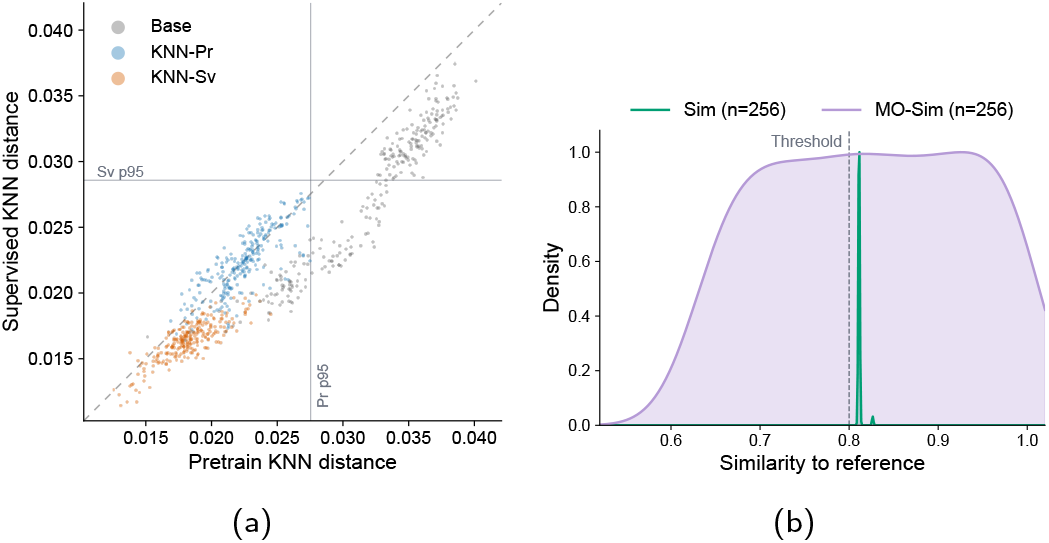
(a) Scatter plot of the KNN distance to the pretraining data (horizontal axis) against the KNN distance to the supervised-training data (vertical axis); the p95 threshold of each reference is shown as a dashed line. (b) Similarity distributions to the reference sequence for Sim (similarity constraint) and MO-Sim (similarity secondary objective); the dashed line marks the constraint threshold (0.80).

Regarding the optimization history (Fig. 5), the Baseline predicted MRL rose sharply up to about generation 100 and then continued to rise gradually up to generation 1000. At the same time, the OOD score also rose sharply, individuals exceeding the threshold appeared, and—while fluctuating—individuals over the threshold became the majority. Under the OOD constraint, in contrast, the predicted MRL rose at the same rate as the Baseline, and although the OOD score also rose, its rise became gradual below the threshold. In addition, under the stronger constraint KNN-Sv, neither the OOD score nor the predicted MRL changed after about generation 200, indicating that optimization converged at an earlier stage.

**Figure 5.**
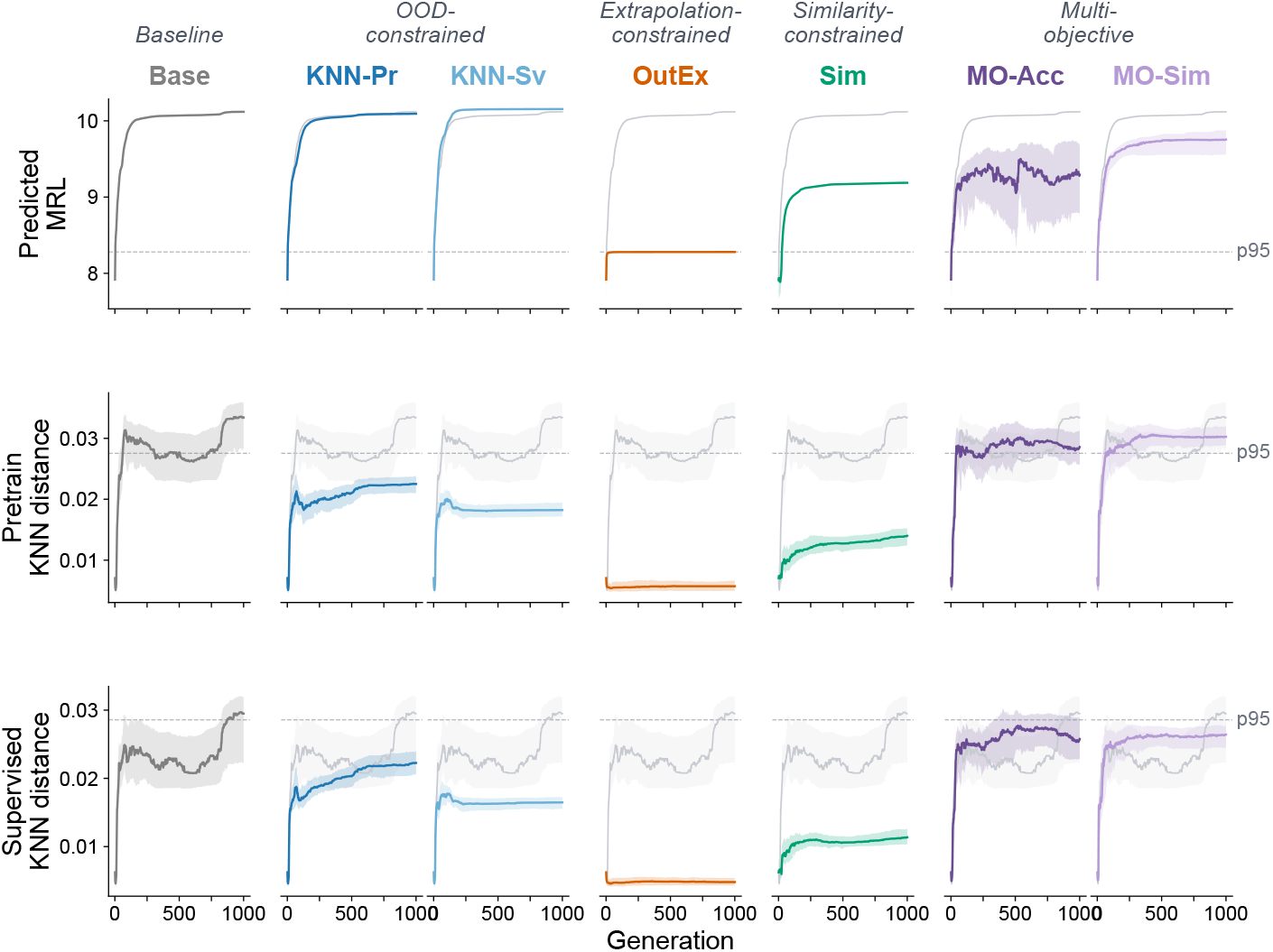
Generational trajectory of the optimization. Top: predicted MRL; middle: pretraining KNN distance; bottom: median (solid line) and interquartile range (band) of the supervised-training KNN distance. Grey is the Baseline reference. Dashed lines mark the p95 threshold.

### Output extrapolation guard

Whereas the OOD constraint attempts to control the optimization space based on the input-side distribution, here we imposed a constraint so that the predicted value stays within the training-data distribution (OutEx) and examined how the search space changes.

Under the OutEx constraint, the final-generation predicted MRL fell entirely below p95, *τ* = 8.28, as designed, with solutions concentrated near this threshold. Furthermore, when we checked the OOD-score distribution under the OutEx constraint, the median OutEx KNN distance (0.0057) was about 1/4 that of KNN-Pr (0.0225) and about 1/3 that of KNN-Sv (0.0182): the OOD score was strongly suppressed, reaching a lower value than under the OOD constraint itself. That is, by holding the output side within the training distribution, the input-side OOD score fell as a side effect. This mirrors the Baseline behavior, in which predicted MRL and the OOD score rise together, and is therefore consistent rather than contradictory. This constraint, however, comes at the cost of plateauing at the MRL ceiling.

### Structural accessibility

Because the structural accessibility around the translation start site affects translation efficiency, we added it as a secondary objective and performed two-objective optimization. As a result, accessibility and predicted MRL are essentially independent (initial population: r= +0.06; after density correction: r *≈ −*0.06), and a trade-off (r =*−*0.57) formed on the Pareto front as a result of the two-objective optimization. The final-generation MRL and OOD score also formed broad distributions. The UMAP likewise shows a tendency to explore more widely than the other conditions. As shown in the UMAP (Fig. 3), because accessibility provides an axis independent of MRL prediction, two-objective optimization is a means of broadening search diversity. The OOD risk, however, is not suppressed. That is, introducing a physical indicator alone does not amount to trust-region control.

### Reference sequence similarity

Taking an arbitrary UTR sequence as the reference, we introduced the similarity to the reference sequence as an indicator. Unlike generic 5′ UTR design, there is a practical demand to modify a sequence starting from a specific reference UTR ((Liu et al., 2025; Chen et al., 2022)). We introduced the reference similarity both as a constraint and as an objective and compared the behavior. When it was a constraint, candidates concentrated at a median predicted MRL of 9.19—about one point below the Baseline—and a population with low OOD risk was obtained. When it was an objective, MRL was broadly distributed over 9–10, and many high-risk sequences with an OOD rate of 82% were included. We also compared the similarity distributions of the two conditions (Fig. 4). When introduced as a constraint, candidates concentrate near the threshold, whereas as an objective the similarity distribution is broad, with many individuals close to a similarity of 1, and identity=1.0 was obtained for 5.1%. When one wishes to search near the reference sequence, introducing it as an objective is considered efficient.

### Threshold sensitivity

Varying the constraint threshold, we compared the median KNN distance of final-generation candidates to the training data (Table 1). Under the KNN-Pr constraint, the tighter the constraint the more the candidate distance shrank (0.020 at p90, 0.023 at p95), while the decrease in predicted MRL was slight. Loosening the threshold to p99, in contrast, gave a median distance of 0.032, nearly indistinguishable from the unconstrained Baseline (0.033), so that the constraint effectively stopped working. The KNN-Sv constraint was similarly threshold-dependent. Under the similarity constraint, the tighter the threshold, the more the distance shrank (0.009 at *θ*=0.90), while the predicted MRL decreased monotonically (9.46 at *θ*=0.70, 8.77 at 0.90).

**Table 1.**
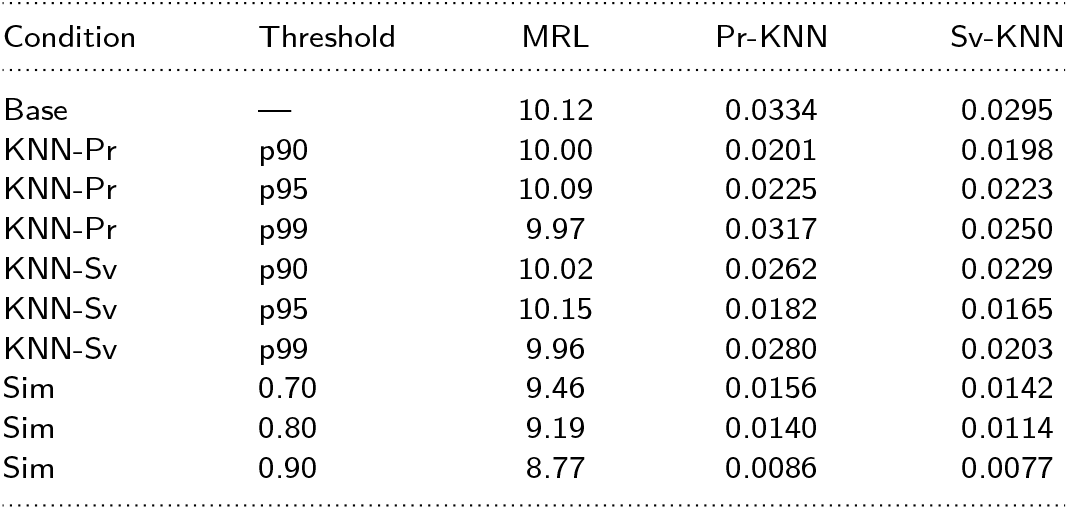
Sensitivity of final-generation candidates to the constraint threshold (seed 42). The MRL column is the median predicted MRL, and the Pr-KNN/Sv-KNN columns are median KNN distances to the training data. The constraint threshold is the feasibility boundary during optimization and is independent of the operational definition of OOD (exceedance of the training-data p95, Section 3.1).

| Condition | Threshold | MRL | Pr-KNN | Sv-KNN |
| --- | --- | --- | --- | --- |
| Base | — | 10.12 | 0.0334 | 0.0295 |
| KNN-Pr | p90 | 10.00 | 0.0201 | 0.0198 |
| KNN-Pr | p95 | 10.09 | 0.0225 | 0.0223 |
| KNN-Pr | p99 | 9.97 | 0.0317 | 0.0250 |
| KNN-Sv | p90 | 10.02 | 0.0262 | 0.0229 |
| KNN-Sv | p95 | 10.15 | 0.0182 | 0.0165 |
| KNN-Sv | p99 | 9.96 | 0.0280 | 0.0203 |
| Sim | 0.70 | 9.46 | 0.0156 | 0.0142 |
| Sim | 0.80 | 9.19 | 0.0140 | 0.0114 |
| Sim | 0.90 | 8.77 | 0.0086 | 0.0077 |

## Discussion

We investigated how to efficiently design low-risk 5′ UTRs that improve translation efficiency, using a high-accuracy predictor and GA optimization. We first aimed to quantify the risk of the designed sequences. The KNN distance in the embedding space of the deep-learning model UTR-Insight (Sun et al., 2022) rose as optimization proceeded (Fig. 5), and was found to represent out-of-distribution better than the pseudo-perplexity PPPL. Lal et al. (2025) related embedding distance to OOD failure in synthetic promoter and enhancer sequences; here we showed the dynamic process by which the KNN distance rises monotonically at each generation of the GA optimization loop as reward hacking proceeds. Our method does not assume that a candidate with a large KNN distance is necessarily mispredicted. Because the predictor’s reported accuracy was evaluated within the training distribution and its validity is not guaranteed outside it, we take as our objective keeping candidates in the region where the validated accuracy applies.

By introducing the KNN distance as a constraint in the genetic algorithm, we showed that low-risk sequences can be explored more abundantly and efficiently while keeping predicted MRL at the same level (Fig. 2). Constraints based on the model’s predicted value or on the similarity to a specific sequence can also suppress OOD risk (Fig. 2). However, a constraint on the predicted value naturally yields a low predicted MRL for the sequences that are finally output; and imposing a similarity constraint can narrow the search space. For the practical demand to modify a sequence starting from a reference, introducing similarity as an objective is more effective than as a constraint: a constraint only guarantees a lower bound (0.80) and concentrates candidates near the threshold, whereas as an objective candidates are distributed continuously up to sequences almost identical to the reference (identity=1.0 for 5.1%) (Fig. 4), allowing fine control of the distance from the reference. When one wishes to generate more diverse candidates, we found it best to perform multi-objective optimization using, in addition to the predicted MRL from the deep-learning model, the similarity to the reference sequence or the structural accessibility as objectives (Fig. 3).

This diversity stems from the orthogonality of accessibility to predicted MRL. Adding accessibility as an objective broadens the search along an axis independent of predicted MRL. That these two parameters are uncorrelated is itself interesting. Given that MRL is widely used as a proxy for translation efficiency, this suggests that UTR-Insight may not sufficiently capture accessibility-derived information in its prediction. Disentangling the causes of this discrepancy with respect to previous studies reporting an association between structure and translation efficiency—including the characteristics of the predictor, the search space, and the proxy nature of MRL—remains a topic for future work.

For the definition of the embedding space, we used the output of the last layer of the encoder block as the readout. Specifically, we used the output of the original UTR-LM encoder block because UTR-Insight reuses the pretrained weights of its predecessor, UTR-LM, while adding a feature transformation module, the ConvTransformerDecoder, to account for local information. In principle, the ConvTransformerDecoder could also have been pretrained, as was the UTR-LM encoder, and its output could therefore have been used as the readout. Had it been pretrained, the convolutional layers might have learned local motif patterns in an unsupervised manner, potentially increasing the resolution of the learned UTR representations and making OOD detection in this representation space more meaningful. This could also be expected to improve MRL prediction performance.

The high correlation in Figure 4 does not necessarily mean that the two overlap spatially. Indeed, in UMAP the two references were largely separated. This correlation is interpreted as arising because both lie in a similar band near the native sequences, so that candidates departing from it move away from both references simultaneously. The reference for the trust region can be set not only from labeled supervised data but also from unlabeled pretraining sequences (native UTRs). Whereas Bayesian optimization, which learns a surrogate on the predicted target value, requires labels, a distance-based OOD constraint can be defined from the sequence embeddings alone, so the region where large and diverse native sequences are distributed can be used as the basis for trust.

In this study, we adopted UTR-Insight as the predictor and a genetic algorithm as the sequence-optimization method. For 5′ UTR design, others have also been proposed, such as UTailoR (Liu et al., 2025), a CNN-based encoder–decoder model, and MOBO-5UTR (Yamada et al., 2025), which combines a DNABERT encoder–decoder with Bayesian optimization in the latent space (LaMBO). These differ in that they all require a generative model (an encoder–decoder) in the optimization loop. Because our method searches directly in discrete sequence space, it requires only an existing predictor, without a generative model, and can incorporate sequence-level hard design constraints—position-fixed motifs (Kozak sequence and splicing motif), sequence length, and prohibitions on GC content, base runs, and start codons—directly and exactly. In latent-space optimization through a decoder, because the candidate sequence depends on the generator’s output distribution, it is generally difficult to impose such positional and combinatorial constraints exactly; and even attempting to secure them by post-hoc filtering is impractical because the generation probability of candidates that simultaneously satisfy multiple conditions is low. In real applications, there are situations where fixing specific positions is essential owing to the requirements of expression vectors and regulatory sequences, and this design flexibility is a practical advantage. A more essential difference lies in the philosophy toward the search space. MOBO-5UTR uses a Gaussian process as the surrogate and incorporates its posterior uncertainty into the acquisition function, working to step actively into uncertain but promising regions and to broaden the design space. Our OOD constraint, in contrast, works to pull candidates back toward the region supported by the training data. The two embody contrasting design philosophies—expansion into unknown regions versus conservation within the trust region—and neither is superior; the choice should be made according to the optimization target and the acceptable risk. Indeed, in-silico comparison of methods has limits, and experimentally testing the proposed sequences of all methods is not efficient either. The choice of method and constraint depends on the understanding of the optimization target.

The threshold sensitivity (Table 1) shows that the threshold itself is a design dial that tunes the trade-off between acceptable risk and MRL, and that its response characteristics differ qualitatively by constraint type. The KNN constraint suppresses OOD efficiently but requires tuning the threshold, whereas the similarity constraint is stable but sacrifices MRL. There is therefore no single right answer as to which constraint to use at which threshold; the designer should choose according to the optimization target and the acceptable risk. That said, the understanding of the optimization target itself is not yet sufficient. It is reported that a compatibility exists between the CDS and the 5′ UTR (Chen et al., 2022), and Verhagen et al. (2026) show that the CDS-leading sequence downstream of the AUG can promote start-codon recognition by the ribosome. 5′ UTR design should therefore preserve the sequence context required for efficient start-codon recognition. Recently, frameworks that co-design the 5′ UTR and the CDS together have been proposed to address this interaction (Liu et al., 2024). Ribosomal selectivity for 5′ UTRs can also vary depending on the species, tissue type, production cell line, and cell cycle. Characterizing such context-dependent effects is therefore critical for successful in-silico sequence optimization. When the properties of the target are not well understood, a practical strategy is to perform optimization under diverse constraints and objectives to broadly explore the search space. Subsequently, once promising design regions have been identified, these regions can be explored more intensively.

The components of our framework—the predictor’s embedding space, the KNN distance to the training data, and its constraint—do not depend on the type of sequence. Therefore, as long as a predictor exists for which an embedding space and a KNN distance can be defined, the same framework should be transferable to other ML-guided sequence design, such as proteins or promoters. Likewise, the OOD constraint works as-is for co-design that includes the CDS.

This study showed that the KNN distance in the embedding space can quantify the OOD risk of ML-guided GA optimization, that using it as a constraint yields candidates within the trust region without sacrificing MRL, and that the extrapolation guard, similarity, and accessibility steer the search space in qualitatively different directions. The choice among these constraints is left to the understanding of the optimization target and the acceptable risk. On that basis, our method does not guarantee the optimal sequence; rather, by keeping candidates within the range where the predicted score is trustworthy, it ensures that subsequent candidate selection stands on trustworthy prediction. In the development of mRNA therapeutics and protein-production cell lines, the experimental cost of synthesizing and evaluating in-silico-designed sequences is large, and testing sequences for which the prediction was wrong is a substantial loss. By focusing on regions where the predictor is reliable, our method increases the proportion of candidates whose predicted performance can be reproduced experimentally. By enabling limited experimental resources to be concentrated on promising sequences, the method has practical value in these applications.

## Author contributions

Ryohei Yamaguchi (Conceptualization [lead], Formal analysis [lead], Investigation [lead], Methodology [lead], Software [lead], Visualization [lead], Writing – original draft [lead], Writing – review & editing [lead]), Chinatsu Mori (Conceptualization [supporting], Methodology [supporting], Writing – review & editing [supporting]), Sota Inoue (Conceptualization [supporting], Methodology [supporting], Writing – review & editing [supporting]).

## Data availability

The datasets underlying this article are available from public resources: Ensembl (Cunningham et al., 2021), the MPRA data of Cao et al. (2021), and GSE114002 (Sample et al., 2019). The architecture and training procedure of the translation-efficiency predictor UTR-Insight are described in Pan et al. (Pan et al., 2025). The source code for the genetic-algorithm optimization and OOD evaluation is not publicly available owing to commercial and intellectual-property restrictions; the Methods section specifies the algorithms, parameters, and metrics in full to enable independent reimplementation.

## Declaration of usage of generative AI and AI-assisted technologies

During the preparation of this work, the authors used Claude-family generative AI models (Opus 4.8, Opus 4.6, Sonnet 4.5, GPT-5.6 Terra) via Claude Code for coding, for drafting figure and table captions, and for translating the Japanese draft into English. The authors take full responsibility for all content of the code and the publication.

